# Proteomic and structural comparison between cilia from primary ciliary dyskinesia patients with a DNAH5 defect

**DOI:** 10.1101/2025.03.14.643267

**Authors:** Charlotte de Ceuninck van Capelle, Leo Luo, Alexander Leitner, Stefan Tschanz, Philipp Latzin, Sebastian Ott, Tobias Herren, Loretta Mueller, Takashi Ishikawa

## Abstract

Primary ciliary dyskinesia (PCD) is a genetic disorder that affects the motile cilia in various organs, leading to recurrent respiratory infections, subfertility, and laterality abnormalities. Traditional diagnostic methods include high-speed video microscopy, immunofluorescence staining, electron microscopy, and genetic screening. However, how different disease-causing variants in the same PCD gene affect clinical presentation, as well as ciliary composition and structure, is not well understood. We investigated the effect of various mutations in DNAH5, an axonemal dynein heavy chain, using mass spectrometry and cryo-electron tomography. We demonstrated differences in the axonemal composition for patients with *dnah5* mutations. Furthermore, we showed that reductions in some ciliary components are patient specific. Some of them, KIAA1430, CFAP97, and DTHD1, were not previously recognized as components of human respiratory motile cilia. Finally, we demonstrated that some differences in protein abundance between wild-type and PCD samples can be observed in the 96-nm repeated unit of the axoneme. Our results suggest that many axonemal components are affected in the DNAH5-defective samples. Additionally, our results highlight that disparate mutations have seemingly distinctive consequences for the axonemal composition.

## Introduction

Primary ciliary dyskinesia (PCD) is a rare genetic disorder, affecting an estimated 1 in every 7500 individuals (Hannah et al., 2022). PCD is characterized by immotility or dysfunction of the motile cilia at various organs of the body (Wallmeier et al., 2020). Motile cilia are hair-like organelles that extend from the cell body and generate a waveform to manipulate the surrounding fluids. Motile cilia consist of the basal body, anchoring the cilium to the cell body, the transition zone, and the axoneme. The axoneme is the core structure responsible for the ciliary motility (Ishikawa, 2017). It is commonly made up of 9 decorated microtubule doublets (MTDs) surrounding two central singlet microtubules. Along the axoneme, the MTDs are decorated with two rows of dynein motor proteins, as well as a variety of structural proteins. Absence or malformation of some of these motor and structural proteins from the axoneme can result in aberrant ciliary function.

People with PCD can display a range of symptoms, including sinopulmonary disease, infertility, cardiac and laterality defects, and hydrocephalus. As there is no definitive diagnostic test for PCD, the European Respiratory Society (ERS) guidelines state that a PCD diagnosis is best obtained using a combinatorial approach (Lucas et al., 2017). Testing procedures for PCD often include nasal nitric oxide measurements, high-speed video microscopy (HSVM), genetic testing, and transmission electron microscopy (TEM) of resin-embedded and stained axonemes. Additionally, immunofluorescence staining (IF) of ciliated cells for commonly affected ciliary components is often performed (Fliegauf et al., 2005).

Axonemal dynein heavy chain 5 (gene: *dnah5*, protein: DNAH5) was identified as a PCD gene in 2002 (Olbrich et al., 2002). Since then, it has become one of the most heavily researched PCD genes, with 15-29% of PCD diagnostic cases attributed to pathological variants in the gene (Zariwala et al., 2007). The *dnah5* gene consists of 79 exons and covers a region of 250 kb (Olbrich et al., 2002), with PCD causal mutations clustering in 5 exons (Hornef et al., 2006). DNAH5 is one of the two heavy chains (HC) making up the outer dynein arm (ODA) complex in humans and corresponds to the γ-HC in *Chlamydomonas reinhardtii*, a popular model organism in cilia motility research (Omran et al., 2000). Loss of DNAH5 from the axoneme has been associated with situs inversus, recurrent respiratory infections, and bronchiectasis (Hornef et al., 2006).

However, how different mutations in the *dnah5* gene affect clinical presentation, and ciliary composition and structure remains unknown. Here, we demonstrate that early, middle, or late truncation of DNAH5 causes similar phenotypes in IF, HSVM, and TEM. To gain insight into the further consequences of DNAH5, we analyzed the composition of the axoneme with mass spectrometry and cryo-electron tomography (cryo-ET). We showed that DNAH5 loss influences motor and structural components throughout the axoneme, such as in the inner dynein arms and the nexin-dynein regulatory complex (N-DRC). Interestingly, several affected proteins (CFAP97, KIAA1439, and VWA3B) had not previously been identified as respiratory motile cilia components. Lastly, we show that DNAH5 truncation is associated with a modest effect on the structure of the 96-nm repeat of the MTD; we observe a loss of density at the site of the N-DRC, further underlining the importance of DNAH5 for ciliary integrity.

## Methods

### Culture of human nasal epithelial cells

The study was approved by the Ethics Committees of the University Children Hospital, Inselspital Bern and of the Canton Bern, Switzerland (reference number 2018-02155). Written informed consent was obtained from all participants or their legal guardians. PCD patient or control cells were obtained by nasal biopsy as described previously (Müller et al., 2021). After the subjects cleaned their nose, nasal epithelial cells (NECs) were obtained by nasal brushings using one interdental brush (IDB-G50 3 mm, Top Caredent; elongated by attaching a 200 μL pipette tip with parafilm) for each nostril. The brush was inserted into and removed out of the inferior turbinate of the nose several times, while rotating and slightly pressing against the lateral edges. Brushes with obtained cells were immediately stored in a RPMI- 1640 media with 20 mM HEPES (cat#R7388, Sigma-Aldrich, Merck, Darmstadt, Germany).

Under aseptic conditions, the elongated brushes were washed with the medium in the tube by pipetting the medium slowly over the brush. Consequently, all cells were removed from the brush by gently pushing and pulling the brush through an adapted 200 μL tip, while keeping the tip submerged in the medium. The tube containing the cells was centrifuged at room temperature (RT) for 5 minutes at 300 rcf. The resulting cell pellet was resuspended in 1.6 mL PneumaCult-Ex Plus basal medium (PC-Ex-Pl, #05040, Stemcell). This fresh cell suspension was then used for various diagnostic tests, such as high-speed video microscopy (HSVM), immunofluorescence (IF), and transmission electron microscopy (TEM). Remaining cells were used for further cell culture.

NECs were initially expanded in T25 and T75 flasks (Sarstedt, Numbrecht, Germany) in PC- Ex-Pl, prepared according to manufacturer’s instructions and supplemented with 96 ng/mL hydrocortisone (# 07925, Stemcell, Vancouver, Canada) and 0.1 mg/mL primocin (ant-pm-05, Invivogen, San Diego, CA, USA). To differentiate the NECs, 2×10^5^ cells were plated on each Transwell polyester membrane 12-well insert (CLS3460, Corning, Corning, NY, USA). An air– liquid interface (ALI) was established by removing the apical media and feeding from the basolateral side. After establishing confluence, the media was switched to PneumaCult-ALI maintenance (PC-ALI, #05001, Stemcell), prepared according to manufacturer’s instructions and supplemented with 480 ng/mL hydrocortisone (# 07925, Stemcell) and 0.1 mg/mL primocin (ant-pm-05, Invivogen), and 4 μg/mL heparin (#07980, Stemcell). Cells at ALI were fed with supplemented PC-ALI medium administered baso-laterally three times weekly. Cells were monitored by light microscopy (Nikon Eclipse TS100, Nikon, Tokyo, Japan). After four weeks at ALI, the cells were heavily ciliated and ready for further processing.

### Whole exome sequencing

Analysis of pathogenic or likely pathogenic variants in PCD-associated genes is performed via next-generation sequencing of the whole exome. For genetic analysis, patients’ blood samples were collected in EDTA tubes and sent to the Molecular and Genetic Diagnostic Lab at the University Hospital Geneva in Geneva, Switzerland or the Department of Human Genetics at the Inselspital in Bern, Switzerland.

### High speed video microscopy

Ciliary motility of freshly harvested or ALI cells was analyzed by HSVM as described previously (Müller et al., 2021). When assessing the motility of freshly acquired ciliated cells from a nasal biopsy, a 60 μL drop of cell suspension was applied to a round 200 μm thick rubber spacer (Ref 64540, Grace Bio-Labs, Bend, OR, USA) on a standard glass slide (631-1553, VWR, Dietikon, Switzerland) and covered tightly with a standard 22×22 mm cover slip (DURAN GROUP, Wertheim, Germany).

When performing HSVM for ciliated ALI cultures, medium from the basolateral chamber was transferred to the apical chamber. Then, the cells were scratched off from the membrane of the insert using a 1000 μL filter tip. Finally, 60 μL media with the cell pieces was applied to the round rubber spacer on a standard glass slide, as described above. Videos were recorded at 300 frames per second at RT on an inverted bright field microscope (Olympus IX73) equipped with a heating plate and a digital CMOS high-speed camera (300 fps, Grasshopper 3 GS3- U3-32S4M-C, USB 3.0; FLIR, Wilsonville, Oregon, USA). The videos were analyzed using the analysis software Cilialyzer (Schneiter et al., 2023) to assess the ciliary beating pattern (CBP) and beating frequency (CBF).

### Immunofluorescence staining

IF analysis was performed as described previously (Müller et al., 2021). The cell culture medium was transferred from the basolateral chamber to the apical chamber of matured ALI cultures. Then, the cells were scratched off the membrane using a 1000 μl pipette tip. Then, the cell mixture was added to a 15 mL Greiner tube and centrifuged at 300 rcf for 5 minutes at RT. After aspiration of the supernatant, the cell pellet was resuspended in 1 mL Accutase (Stemcell, Vancouver, Canada). The tube was placed in the 37 °C incubator for 30 minutes. After the incubation, the cells were gently resuspended and centrifuged at RT for 5 minutes at 300 rcf. After removal of the supernatant, the cell pellet was resuspended in an appropriate amount of RPMI medium (#R8758, Sigma-Aldrich). Finally, 50 μL was added to each slide (631-1553, VWR, Dietikon, Switzerland) and left to air dry.

Cells were fixed using 200 μL of 4% paraformaldehyde for 15 minutes at RT. Afterwards, cells were washed with PBS. The cells were permeabilized with 200 μL permeabilization buffer (PBS with 0.2% triton) for 10 minutes at RT, after which another round of washing with PBS was performed. 200 μL blocking buffer (PBS with 5% milk powder (#T145.2, Carl Roth, Karlsruhe, Germany) was applied to each slide and left to incubate for 1 hour at RT. The primary antibodies were diluted in blocking buffer as per the specifications in Table S1. The blocking buffer was removed from the slides and the slides were lightly dabbed with a paper towel to remove excess buffer. The cells were incubated overnight at 4 °C in a wet and sealed chamber with the primary antibodies. After overnight incubation, an initial short wash, then one 3-minute wash, and finally two 10-minute washes with PBS were performed. 200 μL of each secondary antibody was added to the slides in low light conditions. The slides with secondary antibodies were kept protected from light and at RT for 1.5 hours. Afterwards, the antibody solutions were removed, and the slides were again washed four times with PBS as before. Lastly, the slides were mounted with one drop of Antifade Mounting Medium with DAPI (ENZO Life Sciences, Farmingdale, NY, USA, #ENZ-53003), covered with a 22×22 mm cover glass (Duran Group, 235503207), and stored at 4 °C protected from light until image acquisition.

For staining with DNAH11 and DNAH9 antibodies, an adapted staining protocol was performed: Washing steps were always performed using PBS-Triton. After permeabilization, the cells were washed with 200 μl PBS-Triton three times; once for 3 minutes and two times for 10 minutes. The blocking buffer was prepared with PBS-Triton and the antibodies were diluted in PBS-Triton. Incubation with secondary antibodies was shortened to 30 minutes. Before mounting the cells, we performed one additional washing with PBS. Images were taken with a Leica DMI4000 B Multipurpose Fluorescence System using the x63 oil objective. A detailed list of antibodies including dilutions are provided in Table S1.

### Transmission electron microscopy

Transmission electron microscopy (TEM) was performed as reported previously (Müller et al., 2021). The tissue from fresh nasal brushings and ALI cultures were fixed in 2.5% glutaraldehyde (GA, 350 mOsm, buffered with 0.15 M HEPES at pH7.4) and postfixed with 1% OsO4 (EMS, Hatfield, PA, USA) in 0.1 M Na-cacodylate-buffer (Merck, Darmstadt, Germany) at 4 °C for 1 hour. For ciliated cells grown at ALI, half of a single membrane was cut out of the insert holder using a scalpel and placed in a 1.5 mL Eppendorf tube containing GA.

The fixed tissue was three times washed with 0.05 M maleic NaOH-buffer for 5 min. Afterwards, the tissue was dehydrated with 70%, 80%, and 96% ethanol (Alcosuisse, Rüti b. Büren, Switzerland) for 15 minutes each at RT. Afterwards, the tissue was immersed in 100% ethanol (Merck, Darmstadt, Germany; 3x, 10 min), in acetone (Merck, Darmstadt, Germany; 2x, 10 min) and in acetone-Epon (1:1) overnight at RT. The following day, the tissue was embedded in Epon (Sigma-Aldrich, Buchs, Switzerland) and left to harden at 60 °C for 5 days. The orientation of tissue from ALI inserts was observed under a light microscope. A small block of Epon containing the tissue was subsequently sawed out and realigned to maximize the number of orthogonal cilia cross sections after ultra-thin sectioning.

An ultramicrotome UC6 (Leica Microsystems, Vienna, Austria) was used to produce ultra-thin sections (70–80 nm). The ultra-thin sections were mounted on 200 mesh copper grids (Plano GmbH, Wetzlar, Germany) and stained with a Leica Ultrostainer (Leica Microsystems, Vienna, Austria) using UranyLess (EMS, Hatfield, PA, USA) and lead citrate. Sections were viewed on a transmission electron microscope (Tecnai Spirit, FEI, Brno, Czech Republic) equipped with a 4 Megapixel digital camera (Veleta, Olympus, Soft Imaging System, Münster, Germany) at 80 kV. For each patient, 30 to 50 images at 60,000× magnification were captured providing at least 100–200 orthogonal ciliary transects. Sometimes, single axis tilting of the specimen holder up to 40° was used to increase the delineation of axonemal structures. The axonemal structures were systematically evaluated and scored according to the international consensus guidelines on TEM in PCD diagnosis (Shoemark et al., 2020).

### Cilia isolation for cryo-ET and MS

Isolation of human respiratory cilia grown at ALI was performed as previously described (Lin et al., 2014). After removal of the culture media from the basolateral chamber, the cell culture inserts were washed twice with RT PBS for 5 minutes. After ice-cold PBS was added to both the apical and the basal compartments, the plate was placed on ice for 5 minutes. The ice-cold PBS was removed and 100 μl deciliation buffer (DC buffer, 10 mM Tris pH 7.5, 50 mM NaCl, 10 mM CaCl2, 1 mM EDTA, 0.1% Triton X-100, 7 mM β-mercaptoethanol, 1% protease inhibitor cocktail (P8340, Sigma-Aldrich)) was added to the apical side of each insert and left to incubate on ice for 2 minutes. The apical solution containing the isolated cilia was collected, transferred to a pre-cooled microcentrifuge tube and centrifuged at 4 ℃ at 500 rcf for 1 minute. The supernatant was placed into a new pre-cooled microcentrifuge tube and centrifuged at 4 ℃ at 5000 rcf for 5 minutes to pellet ciliary axonemes. After removal of the supernatant, the ciliary pellet was gently dispersed in the resuspension buffer (30 mM HEPES pH 7.3, 1 mM EGTA, 5 mM MgSO4, 0.1 mM EDTA, 25 mM NaCl, 1 mM dithiothreitol, 1% protease inhibitor cocktail (P8340, Sigma-Aldrich) and 100 g/ml soybean trypsin inhibitor (T9128, Sigma-Aldrich, Merck, Darmstadt, Germany) and kept on ice until further processing.

### Cryo-electron tomography

R3.5/1 Holey carbon copper grids (QUANTIFOIL, Großlöbichau, Germany) were glow-discharged under UV light for 10 minutes. 3 μl of 10 nm gold bead solution (752584, Merck, Darmstadt, Germany) was applied to each grid as fiducial markers. The axoneme suspension was brought to 0.1-0.4 mg/ml protein concentration before application to the grids. We performed manual back side blotting for 3 seconds (Nr.1, Whatman, Maidstone, UK). The grids were plunge-frozen by the Cryoplunge3 (Gatan, Pleasanton, CA, USA) with >80% humidity in the humidity chamber. The tilt series were acquired on a 300 kV Titan Krios (Thermo Fisher, Waltham, MA, USA) with a K2 camera and GIF-Quantum energy filter (Gatan, Pleasanton, CA, USA). Tilt series from −60° to 60° were collected with a 2° increment using a bidirectional tilt scheme starting from 0° using SerialEM [Mastronarde, 2005]. The total electron dose used for both datasets was around 60-80 electrons per Å^2^. The frames of the dose fractionated normalized micrographs were aligned using the IMOD *alignframes* command. Tomograms were reconstructed using IMOD [Kremer et al, 1996]. CTF correction was performed using the CTFplotter in IMOD.

### Subtomogram averaging

Particle picking and initial subtomogram averaging was performed as described previously (Table S2) (Bui and Ishikawa, 2013; Zimmermann et al., 2023). Microtubule doublets (MTDs) were manually picked in IMOD using the *3dmod* viewer (Kremer et al., 1996). We made use of an in-house developed program called axoneme_aln to determine particle position and orientation. Briefly, points along one MTD were interpolated at a 24-nm distance from each other. These subvolumes (160×160×160 pixels at a pixel size of 8.68 Å) were aligned along one MTDs, under the assumption that they have similar Euler angles. The 24-nm subtomograms of all nine MTDs were then aligned based on (constraint) cross-correlation with a reference during several alignment rounds, leveraging the pseudo nine-fold symmetry of the axoneme. Afterwards, aligned particle positions and orientations were converted into a RELION-readable star file. Reference-free 3D classification and refinement was performed in RELION (Scheres, 2012). All the previously aligned subtomograms (at 24-nm periodicity) were extracted and binned to 17.36 Å/pixel. The 3D Autorefine function was used to optimize the alignment of the subtomograms. Then, reference-free 3D classification, without alignment, with a mask of a single central outer dynein arm (ODA) was carried out in RELION. The outputs of the ODA classification were finally refined. We used the IMOD software (Kremer et al., 1996) and UCSF ChimeraX (Meng et al., 2023) for visualization of the tomographic slices and isosurface renderings, respectively.

### Mass spectrometry

Wild-type and PCD patient axonemes were isolated from the respiratory cells grown at ALI according to the cilia isolation protocol described above. After incubation with DC buffer (10 mM Tris pH 7.5, 50 mM NaCl, 10 mM CaCl2, 1 mM EDTA, 0.1% Triton X-100, 7 mM β-mercaptoethanol, 1% protease inhibitor cocktail (P8340, Sigma-Aldrich, Merck, Darmstadt, Germany)) on the insert, the apical solution containing the axonemes was transferred to a 1.5 mL Eppendorf tube and centrifuged at 4 ℃ at 5000 rcf for 5 minutes. The supernatant was discarded. The axonemal pellet was snap-frozen in liquid nitrogen and stored at −80 ℃ until use. Further processing was performed according to a previously published protocol (Leitner et al., 2014), with omission of the crosslinking steps. Briefly, the samples were denatured using urea, alkylated with iodoacetamide and trypsin digested. The digested samples were purified using reverse-phase solid extraction, evaporated and redissolved in the MS mobile phase A. The isolated cilia samples were analyzed by liquid chromatography-tandem mass spectrometry (LC-MS) in the data-dependent acquisition mode on an Easy nLC-1200 liquid chromatography instrument couple to an Orbitrap Fusion Lumos mass spectrometer (ThermoFisher Scientific). Raw spectral data was analyzed by MaxQuant (Cox and Mann, 2008). Quantitative analysis was performed in Perseus (Tyanova et al., 2016). During data processing in Perseus, common contaminants were removed, entries with fewer than 2 valid values were removed, and missing values were imputed and log-transformed. The resulting values were normalized by subtracting the median value for each sample to account for differences in protein amounts between samples. Then, differentially present proteins between patient and healthy volunteer samples were identified using two-sided T-testing. Network analysis was performed based on reported interactions in the STRING database v12 (Szklarczyk et al., 2023). The heat maps were generated in Python. MS intensities were log-transformed and normalized to the TUBB4B content of the sample before plotting. A full overview of software used in this study can be found in Table 1.

**Table 1.**
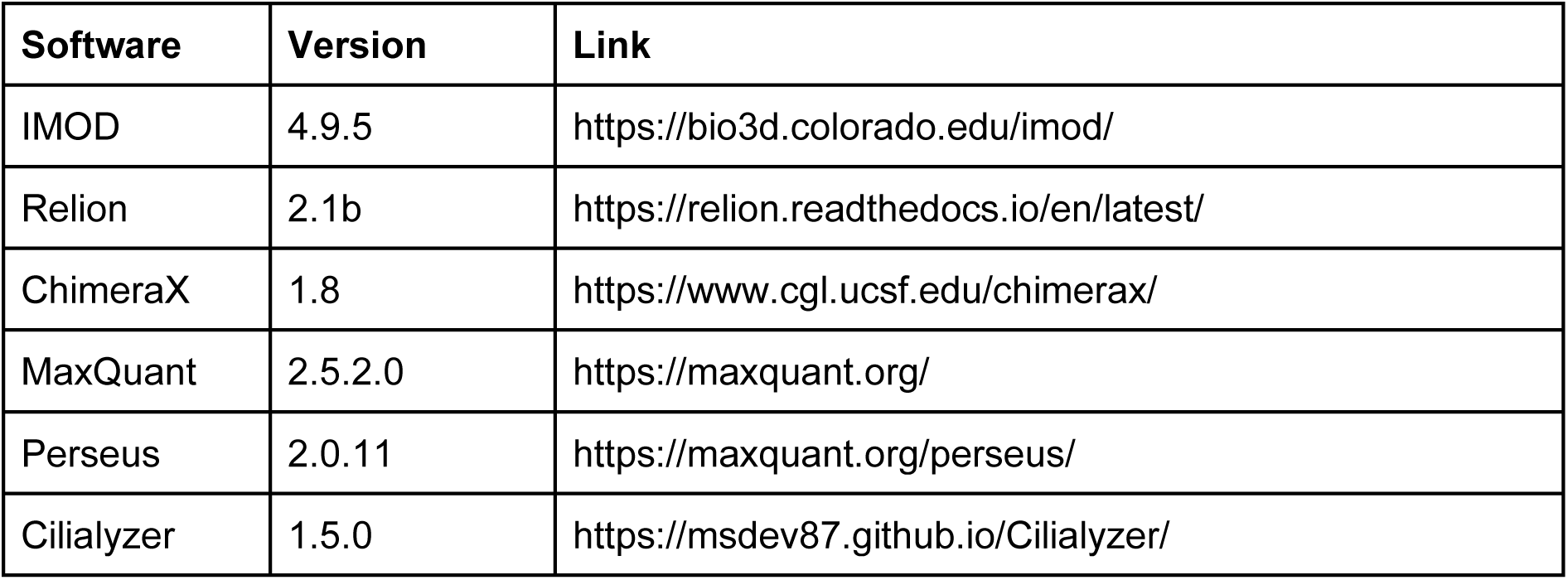
Software used in this study.

| Software | Version | Link |
| --- | --- | --- |
| IMOD | 4.9.5 | https://bio3d.colorado.edu/imod/ |
| Relion | 2.1b | https://relion.readthedocs.io/en/latest/ |
| ChimeraX | 1.8 | https://www.cgl.ucsf.edu/chimerax/ |
| MaxQuant | 2.5.2.0 | https://maxquant.org/ |
| Perseus | 2.0.11 | https://maxquant.org/perseus/ |
| Cilialyzer | 1.5.0 | https://msdev87.github.io/Cilialyzer/ |

## Results

### Disparate mutations in the ODA γ-HC show similar phenotypes in IF and TEM

The genetic mutations concerning the outer dynein arm (ODA) γ-HC *dnah5* gene for each patient in this study are outlined in Table S3. Patient PD122 suffers from a large deletion in chromosome 5, a syndrome known as Cri-du-chat (Rodríguez-Caballero et al., 2010), including the genomic region that contains the *dnah5* gene. Additionally, the patient has a single nucleotide variant (4348C>T) in the gene that results in a premature stop at Gln1450 (rs771663107). In high-speed video microscopy (HSVM), PD122 cilia demonstrated immotility (data not shown). Consistent with the large deletion in chromosome 5 and the loss of function, patient PD122 was diagnosed at a relatively young age (Kuehni et al., 2010), suffers from serious respiratory illness and situs inversus totalis (Table S4). PD122 was classified as an ERS class 1 hallmark defect in transmission electron microscopy (TEM) analysis (Shoemark et al., 2020), supporting a PCD diagnosis for these patients on the basis of ODA defects (Figure 1b). The majority of the axonemata showed ODA defects in both the proximal and distal regions. In ∼34% of axonemata of cells grown at ALI, ODA were present. Additionally, defects in inner dynein arms (IDAs) were also penetrant (Figure 1b), as they were present in ∼55% of axonemata for this patient. As part of the diagnosis procedure, we performed IF staining with a panel of PCD-related proteins, and found that DNAH5 (Figure 1c), dynein axonemal intermediate chain 1 (DNAI1), DNAI2, DNAH11, DNAH9 (data not shown) were missing in the cilia of this patient. Other PCD-related proteins, such as growth arrest specific 8 (GAS8), coiled-coil domain containing 40 (CCDC40), and dynein axonemal light intermediate chain 1 (DNALI1) were present (data not shown).

**Figure 1.**
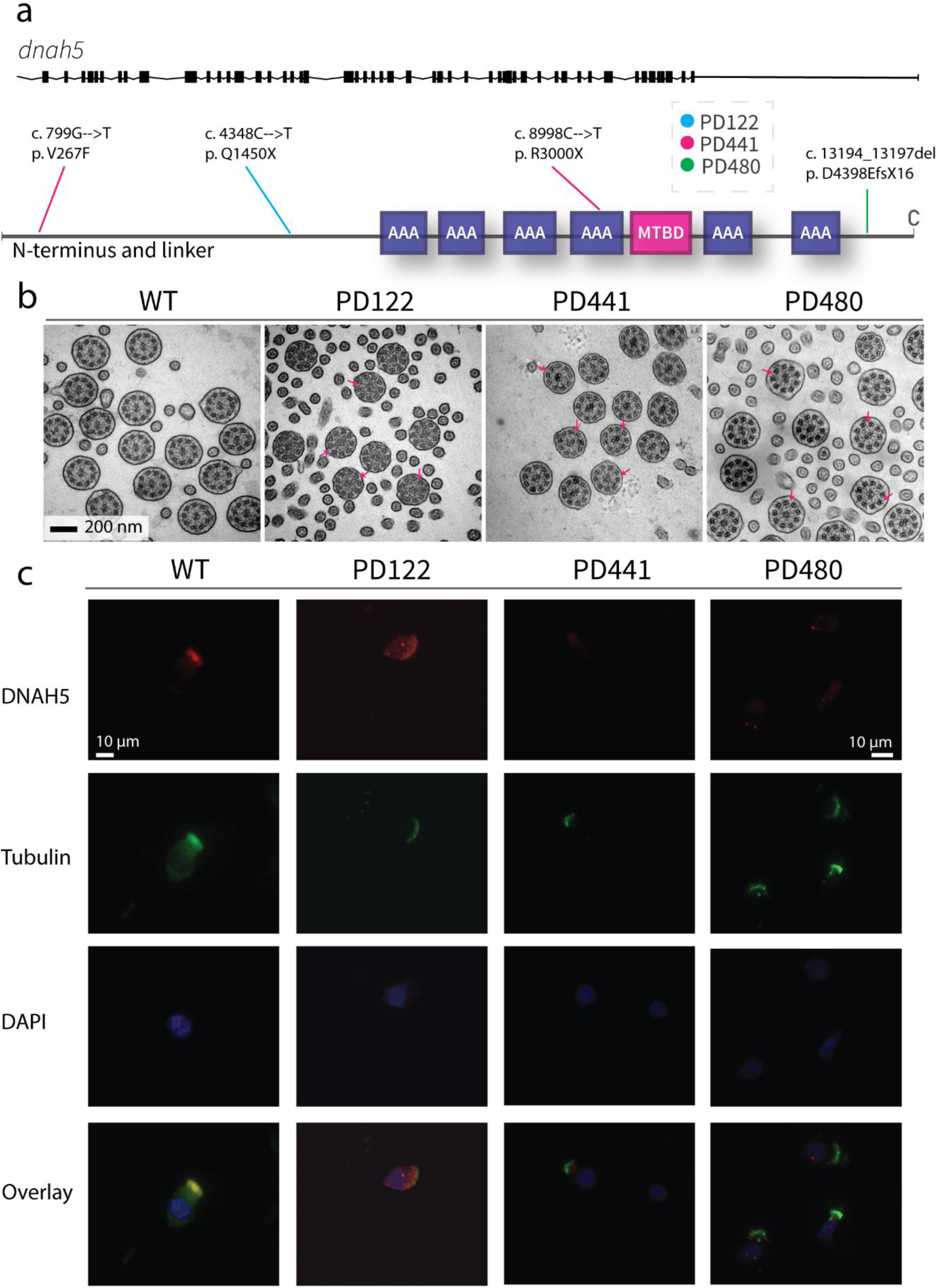
Traditional diagnostic procedures for all patients demonstrate similar phenotypes in TEM and IF. A. Schematic of the genetic variants in *dnah5* of patients included in this study. B. Representative images of the cross section of wild type and patient cilia as imaged by transmission electron microscopy. Extensive loss of ODA can be seen in all patient samples. Pink arrows indicate examples of clearly visible ODA loss in patient samples. C. Representative immunofluorescence images of wild type and PCD patient cilia. All patients show a loss of DNAH5 (red) from the axoneme. Tubulin is stained in green and DNA (DAPI) in blue.

The second patient, patient PD441, was shown to carry a heterozygous nonsense mutation (8998C>T) resulting in a termination at Arg3000 of DNAH5. Additionally, this patient carries a heterozygous missense mutation (799G>T) resulting in a phenylalanine instead of a valine at the protein level at position 267. This change on the first nucleotide of exon 7 is predicted to be likely pathogenic. HSVM showed PD441 to have immotile cilia (data not shown). PD441 presented neonatal respiratory distress in early life, a feature of PCD (Mullowney et al., 2014). Additionally, PD441 suffers from chronic rhinosinusitis, cough, and recurrent lung infections (Table S5). Like PD122, PD441 was shown to possess a ERS class 1 hallmark defect; ODA defects were detected in ∼87% of axonemata with no significant proximo-distal preference (Figure 1b). A modest IDA defect (40%) was observed in PD441 axonemata. DNAH5 was shown to be missing from the axoneme in roughly half of the imaged ciliated nasal cells for this patient (Figure 1c). Interestingly, we observed proximally reduced DNAH5 in the remaining ciliated cells. Additionally, in IF staining of ciliated cells grown at ALI, DNAH9, DNAH11, DNAI1, and DNAI2 were shown to be missing from the axoneme.

In whole exome sequencing (WES), patient PD480 was shown to carry two deletions; a small deletion at the C-terminal domain, predicted to result in a frame shift and a subsequent premature termination in exon 76 (rs727502971), and a larger heterozygous deletion in exon 69. PD480 was found to have immotile cilia in our analysis (data not shown). Consequently, PD480 suffers from chronic cough and bronchiectasis (Table S6). Similar to PD441, PD480 showed an almost complete ODA defect with no proximo-distal preference (91%, figure 1b).

Additionally, a reasonable IDA defect was detected with a penetrance of 65%. Similar to in the other patients, we found that DNAH5 (Figure 1c), DNAH9, DNAH11, DNAI1, and DNAI2 were missing in IF. GAS8 and DNALI1 were found to be present.

### Mutations in *dnah5* affect the axonemal composition outside of the ODA complex

To further understand the consequences of *dnah5* mutations on the axoneme, we performed bottom-up label-free quantitative proteomics. Isolated and demembranated axonemes from the DNAH5-defective PD441 and PD480, and healthy controls (three replicates for each) were denatured, digested, and purified before LC-MS data acquisition. We then compared the intensities for significantly differentially abundant proteins between the samples. We observed that a lack of DNAH5 is robustly identified in PD441 and PD480 samples (Figure 2a, Figure S1), with it being the most striking hit in terms of fold change and significance. Additionally, other ODA component proteins, such as β-HCs DNAH9 and DNAH11, and ICs DNAI1 and DNAI2, were significantly reduced in the sample. We have highlighted the names of these ODA proteins in the volcano plot summarizing the MS results (Figure 2a).

**Figure 2.**
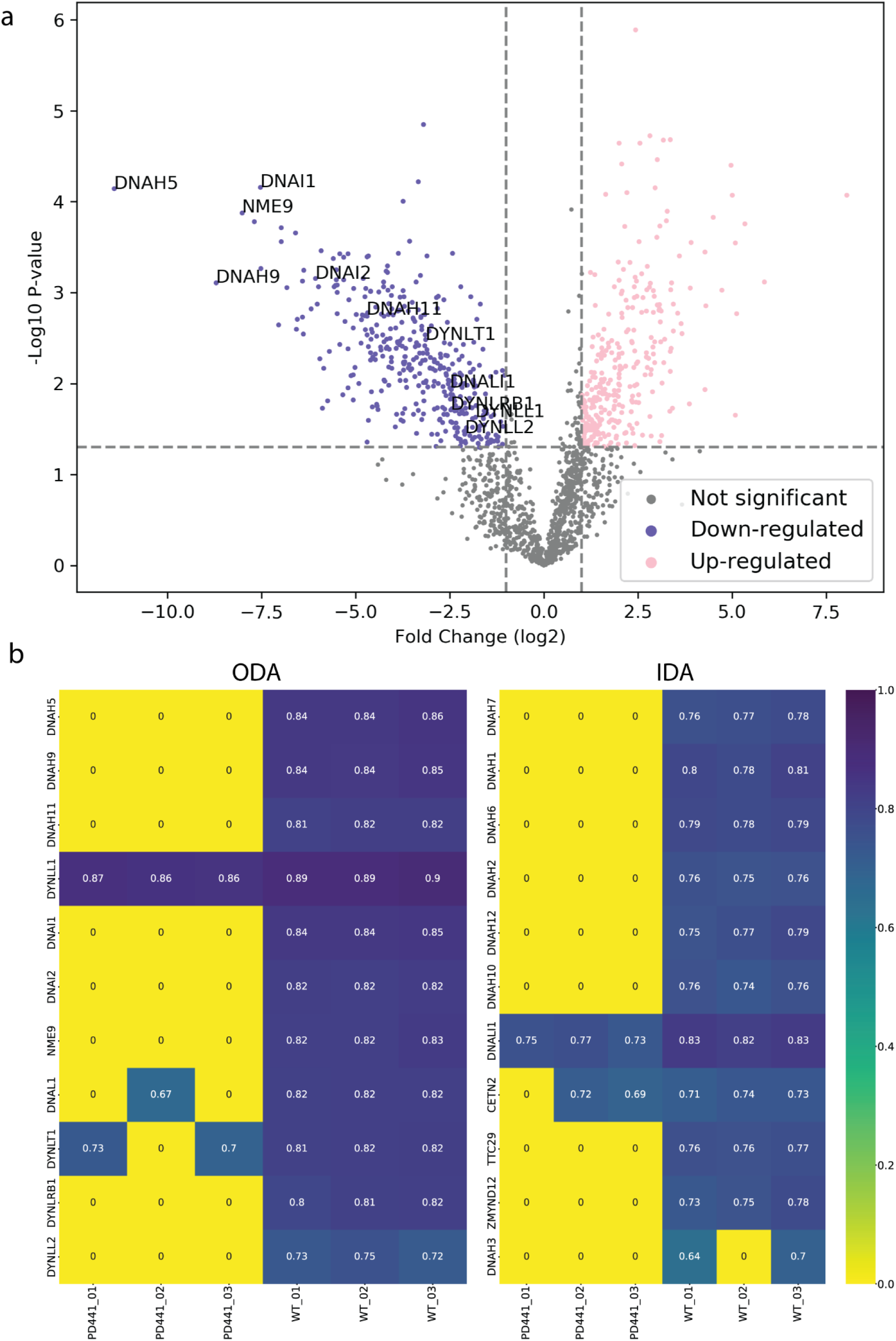
Loss of DNAH5 from the axoneme is associated with loss of other axonemal components. A. Volcano plot of the significantly less (blue) and more (pink) abundant ciliary proteins inPD441 compared to WT. ODA components are highlighted by their gene names. B. Heat maps comparing the intensities of ODA and IDA components between wild type and PD441 isolated cilia. All samples were normalized to their respective sample TUBB4B intensity, although the numbers are not directly proportional to the real protein amounts.

Interestingly, in the PCD patients, we observed a reduction in heavy chains belonging to IDAs (Figure 2b). We also observed a reduced abundance in proteins belonging to other classes of axonemal components, such as the microtubule inner proteins (MIPs), nexin-dynein regulatory complex (N-DRC), and microtubule inner proteins (MIPs) (Figure S1).

### DNAH5 truncations differently affect axonemal protein composition

We compared the protein composition for PD441 and PD480 to understand how early or late truncations of DNAH5 affect axonemal protein composition. We identified several key differences in protein composition between the two patient samples. Firstly, glutathione S-transferase A2 (GSTA2) was lost from PD441 axonemes, but present at wild type levels in PD480. Additionally, several components belonging to two complexes were missing from PD441 axonemes (Figure 3a), compared to PD480 axonemes. Decreased IFT components such as IFT-140 and IFT-88 are part of the IFT-A and IFT-B complex, respectively (Lacey et al., 2023). The IFT-A and IFT-B complexes are essential for intraflagellar transport (IFT), the process required to build ciliary axonemes. Furthermore, the proteomics analysis for PD441 and PD480 indicated a reduction of several proteins that were not earlier recognized as associated with motile respiratory cilia: Cilia and flagella associated protein 97 (CFAP97, also called KIAA1430), Von Willebrand factor A domain-containing 3B (VWA3B), and death domain containing 1 (DTHD1) (Figure 3b). Loss of these proteins was not observed in isolated axonemes from PD480. These proteins are associated with ciliary components in the STRING interaction database (Figure 3c).

**Figure 3.**
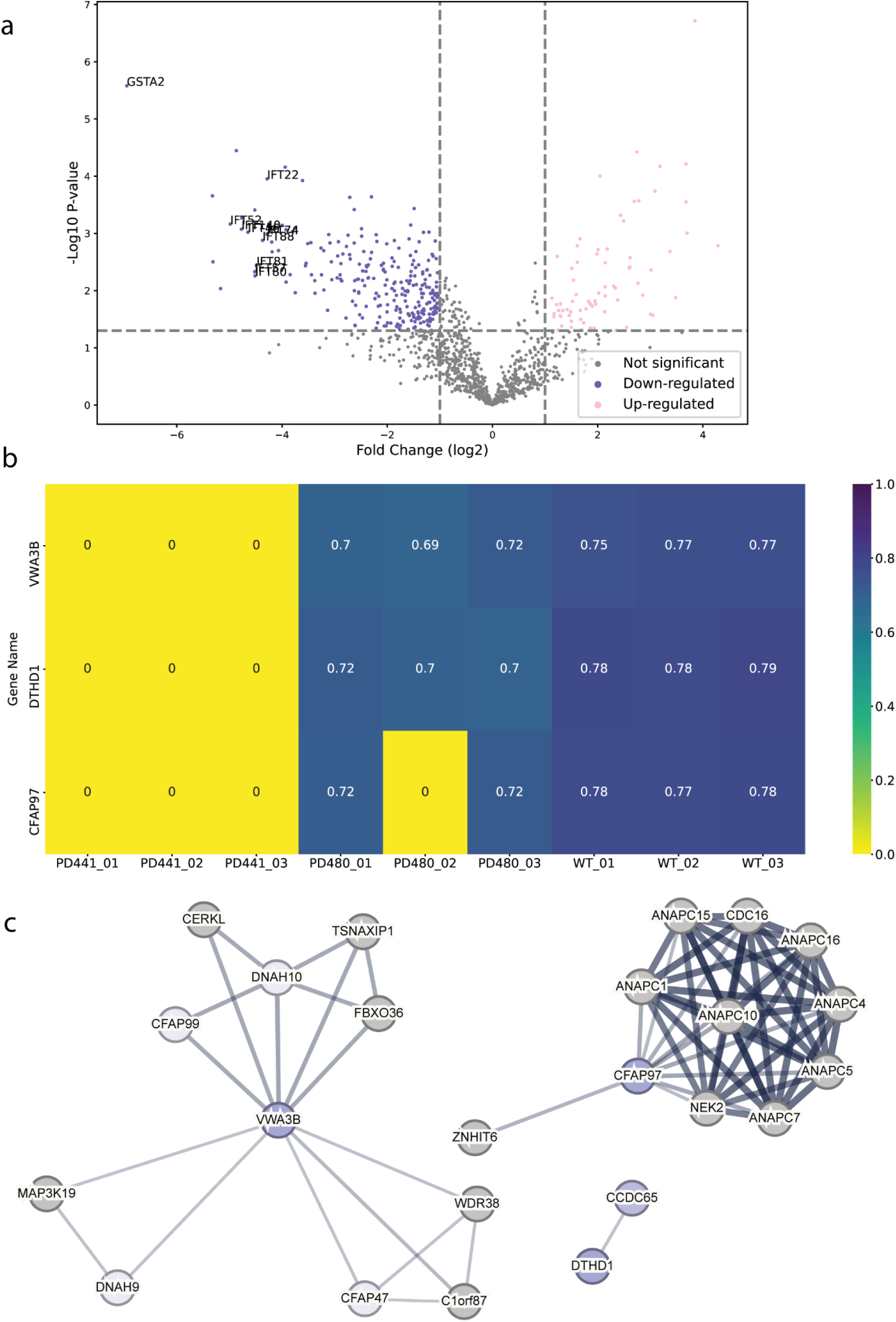
DNAH5-defective patient cilia carrying different dnah5 mutations show distinctive protein composition patterns. A. Volcano plot comparing the significantly less (blue) and more (pink) abundant ciliary proteins in PD441 than in PD480. GSTA2 and several IFT proteins are highlighted by their gene name. B. Heatmap of proteins that are identified in MS of wild type and PD480 cilia, but not in the PD441 samples, suggesting some patient specific differences in the axonemal composition. All samples were normalized to their respective TUBB4B intensity. C. Protein interaction networks for the newly ciliary identified proteins based on reported STRING interactions.

### DNAH5-defective PCD patients with different mutations in *dnah5* show similar structures in cryo-ET

To compare MTD structures in axonemes of PCD patients with a *dnah5* mutation, we performed cryo-ET. We isolated axonemes from control and patient ciliated cells. Tilt images were acquired and reconstructed into 3D volumes. From the 3D volume, we extracted sub volumes with our axonemal structure of interest. These subtomograms were aligned and averaged to obtain a higher signal-to-noise ratio. In the wild type axonemes, the ODAs were arranged along the doublet with a 24-nm periodicity (Figure 4a/b). Both heavy chains were easily identifiable (Figure 4b/c). In the averaged structures from all the patients, we observed a loss of the ODAs from the axonemes (Figure 4d). The only remaining identifiable structure at the location of the ODA is the docking complex (ODA-DC). In addition, we observed rather globular densities between the DCs and towards the external side of the MTD (Figure 4d). When fitting the previously established atomic map of the bovine docking complex (7RRO) (Gui et al., 2021) into our tomographic volume, we observed considerable consistency between the atomic map and our subtomogram average (Figure 4e).

**Figure 4.**
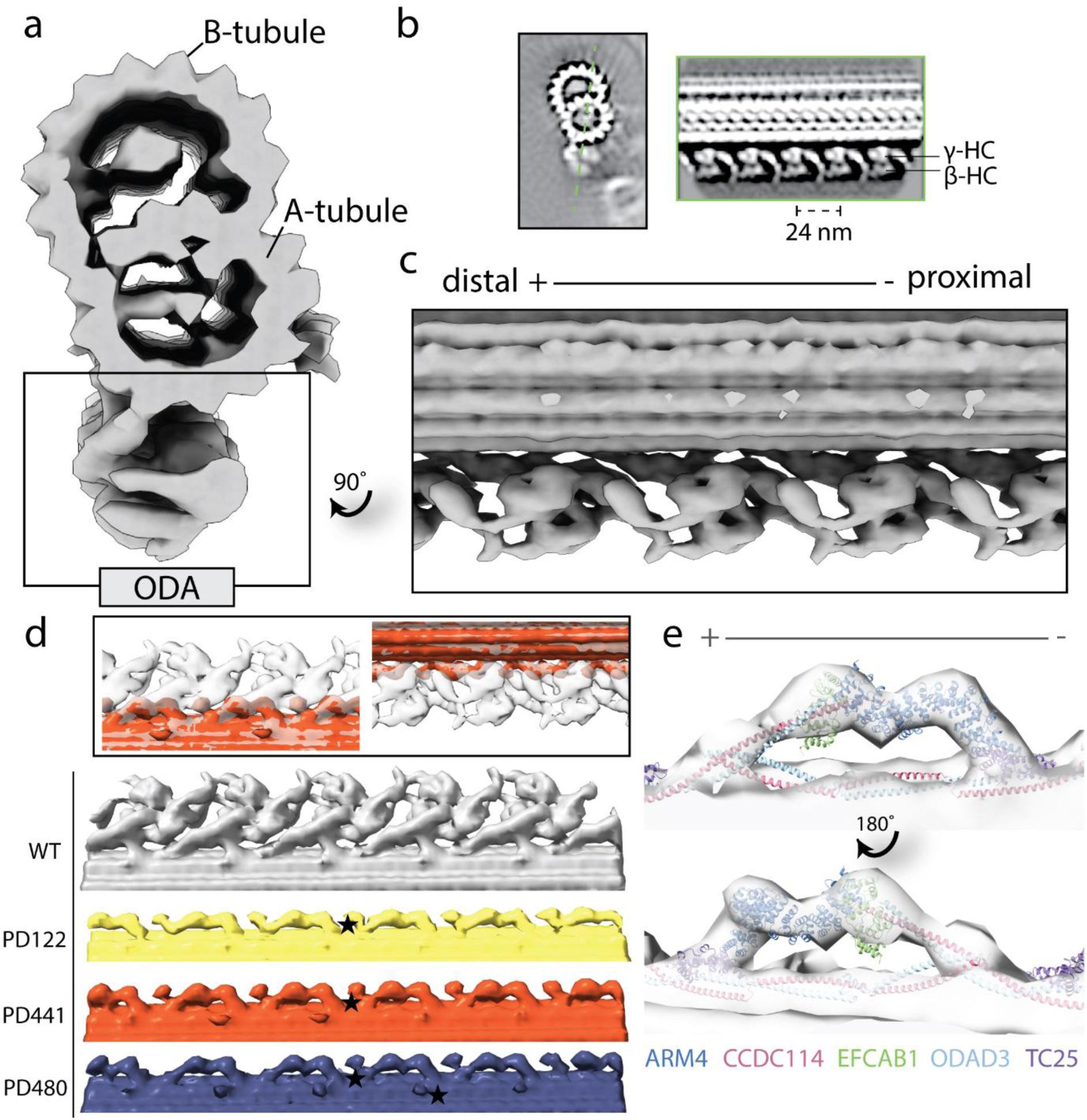
Structural comparison of wild-type and patient outer dynein arms (ODAs) by cryo-electron tomography and subtomogram averaging. A. Isosurface rendering of a wild-type microtubule doublet (MTD) average. The ODAs are highlighted on the bottom of the doublet. B. Tomographic slices of the healthy MTD (left) and the double-headed ODA (right). The orientation of the ODA slice is indicated by the dotted green line. C. Isosurface rendering of wild type MTD with 24-nm periodicity with ODA view. D. Isosurface rendering comparison of wild type and patient MTD with 24-nm periodicity with ODA view. Star denotes an example of the additional globular density identified in PCD patient axonemes. E. Fitting of the ODA docking complex chains from atomic model RRO7 (Gui et al., 2021) into the tomographic map of PD480.

We compared the structures of the 96-nm periodic unit for all patients in this study and WT. In the consensus structure, all patients lack ODAs and are just left with the DCs (data not shown). When comparing the 96-nm periodic structure for PD441 and WT, we observed several locations with a reduction in or lack of density (Figure 5). In the consensus map, we see a minor reduction in density in one of the IDAs, namely DNAH7, which is located at the position of IDAe of Chlamydomonas (Figure 5a). This corresponds to a reduction in DNAH7 in these samples, which was identified in MS (Figure 2b). However, further investigation did not show this reduction with reproducibility. We also observed a reduced density at the site of one of the dimeric inner dynein HCs in PD441 (Figure 5b/d). This corresponds to DNAH10, which was found to be below the detection threshold by MS.

**Figure 5.**
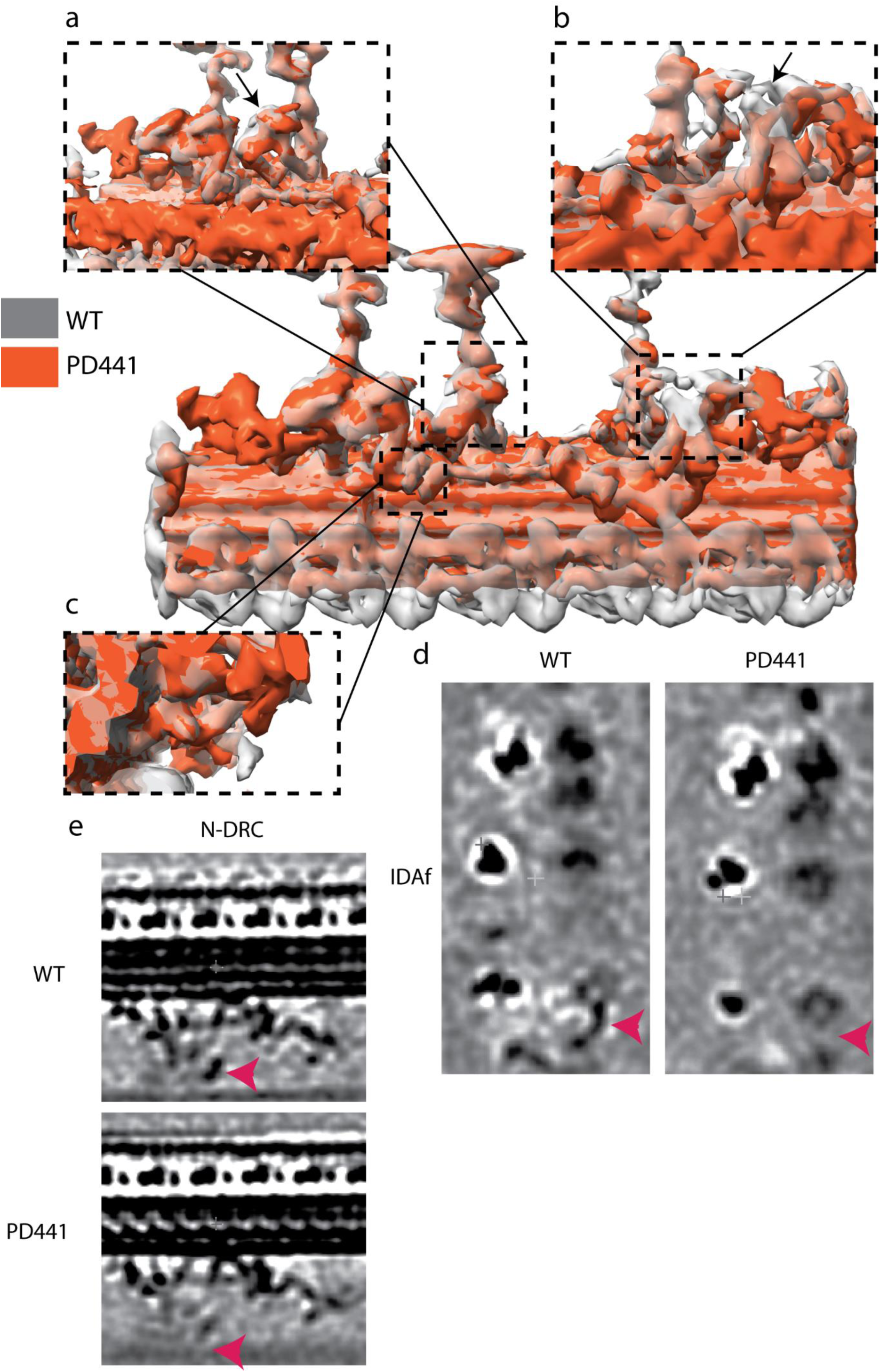
Overlay of 96-nm periodic structure of the wild-type (grey, transparent) and PD441 (orange) microtubule doublet. Zoomed in view of several components that are demonstrating reduced density in PD441 compared to WT: A. IDAe (DNAH7). B. IDAf heavy chain a (DNAH10) (both shown by black arrows). C. N-DRC. Tomographic slices of the reduced D. IDAf, and E. N-DRC structures, generated with *3dmod*. Pink arrows in d and e denote the location of the IDAf HC and the N-DRC, respectively.

Finally, we did not observe any density at the site of the portion of the N-DRC approaching the adjacent MTD in PD441 (Figure 5c/e). This loss can also be seen in the proteomics data, as many N-DRC components were undetected in MS measurements (Figure S1). MIPs, which were found to be reduced in PCD samples in MS, were found to be present in comparable densities between WT and PD441 (Figure S2b), though a general loss of TEKTINs can be observed in the structural analysis of both samples.

## Discussion

In this study, we found that disparate mutations in *dnah5* lead to highly similar phenotypes in classical PCD diagnostic methods, including IF, TEM, HSVM. However, we found that differences in the axonemal composition can be observed by MS for patients with distinct *dnah5* mutations. Our findings suggest that earlier truncations in DNAH5 may result in a more substantial reduction in the axonemal components compared to a later truncation in DNAH5. Additionally, our cryo-ET analysis confirms the loss of ODA from the axoneme and suggests that changes in protein abundance in the axoneme can be identified at a structural level.

### DNAH5 loss of function corresponds to a lack of ODAs in the axoneme

Through WES analysis of three PCD patients, we identified two variants that were previously described in the literature: rs771663107 (Failly et al., 2009; Ferkol et al., 2013) in PD122 and rs727502971 (Hornef et al., 2006) in PD480. The small deletion gene variant rs727502971 has been previously described in two individuals from Switzerland (Hornef et al., 2006). Similarly to PD480, the afflicted individuals suffered from completely immotile cilia. We found that, regardless of the position of the DNAH5 truncation, ODA heavy chains are lost from the axoneme. We observed that DNAH5 is instead accumulated in punctae in the cell or present near the basal body. Additionally, we saw that the other ODA heavy chains, DNAH9 and DNAH11, are weakly and diffusely present throughout the cytoplasm. This is consistent with findings in a previous study (Fliegauf et al., 2005), where the authors observed an accumulation of DNAH5 around the proximal perinuclear region and the microtubule-organizing centers. They also observed weak and diffused presence of DNAH9, throughout the cytoplasm in DNAH5-defective cells. However, this presents a different situation compared to *Chlamydomonas reinhardtii*. In *Chlamydomonas*, a HC-γ truncation that preserves the N-terminal tail (*oda2-t*) does not lead to a total loss of ODA from the axoneme (Liu et al., 2008). This suggests that pre-assembly and transport of ODA complexes could be governed by different interactions between organisms. Additionally, we observed in one IF staining that PD441 maintains reduced levels of DNAH5 in the proximal region of some axonemes (data not shown). We did not ever observe such staining patterns for β-HCs DNAH9 or DNAH11, indicating that any presence of DNAH5 does not signify a complete ODA. ODAs were also observed in a minority of axonemata analyzed with TEM (data not shown) but did not represent any substantial class in cryo-ET classification analysis. It is possible that the stability of the axonemes is compromised in isolated axonemes with an DNAH5 defect, and such rare ODAs are not maintained in experimental preparation. Another interesting observation is that even though the patients present similarly in traditional diagnostic methods, clinical manifestations can differ significantly. For example, PD122 suffers from situs inversus totalis (Table S4), whereas PD441 does not have any situs abnormalities (Table S5).

### Earlier DNAH5 truncation is associated with a more extensive reduction in axonemal components

Our MS results indicate that loss of DNAH5 has more extensive effects on the doublet structure than previously believed, with reductions in components belonging to the IDAs and N-DRC among others (Figure S1). *dnah5* mutations are often associated with both outer and inner arm defects (Failly et al., 2009). However, we suggest that many classes of axonemal components are affected in the DNAH5-defective samples. Further research should focus on whether lack of ODAs affect assembly of ciliary components or axonemal stability. Additionally, our results highlight that disparate mutations have seemingly distinctive consequences for the axonemal composition. GSTA2 was not detected in the isolated axonemes of PD441 (Figure 3A). Surprisingly, it was robustly detected in PD480 (Figure S1). GSTA2 was previously found to be highly expressed in respiratory epithelial cells and shown to localize to the ciliary axoneme (Koenitzer et al., 2024). Interestingly, transcriptomics analyses by Koenitzer and colleagues has shown that GSTA2 expression is increased in PCD ciliated airway cells (Koenitzer et al., 2024). Of course, the relationship between mRNA levels and protein abundance is difficult to interrogate (Zhang et al., 2014, 2016). However, these results do indicate that not every *dnah5* variant is created equally, as they might affect GSTA2 localization patterns differently.

We see considerable consistency between the proteomics and cryo-ET data. For example, we saw a decrease in the presence of ODA-DC components, ODAD1/2/3/4, in PD441. In PD480 axonemes, ODA-DC components were found to be present nearly at wild type levels (Figure S1). When we classified the ODA particles in cryo-ET, one of the major classes obtained represented a fraction of the particles where the ODA-DC is completely lost from the MTD of PD441 (∼10% of particles, figure S2a). We did not observe any similar class in PD480 classifications.

### Abundance of proteins that were previously not associated with respiratory motile cilia are reduced in DNAH5-mutant samples

We also discovered a disparity in the ciliary presence of several proteins, namely KIAA1430, CFAP97, and DTHD1, between wild-type and patient samples (Figure 3b). KIAA1430 is conserved in species with motile cilia. Moreover, it was shown that *Drosophila* KIAA1430 orthologue *hemingway* is essential for sperm flagellar assembly and maintenance (Soulavie et al., 2014). Other research localized KIAA1430 to the primary ciliary axoneme (Gupta et al., 2015)]. KIAA1430 has been shown to interact with cell division cycle 16 (CDC16), and several Anaphase-promoting complex (APC) subunits (Figure 3c) (Hein et al., 2015). The APC complex has been shown to regulate cilia length and disassembly in primary cilia (Wang et al., 2014), whereas in motile cilia it has been implicated in regulating cilia polarity (Ganner et al., 2009). Another relatively under-researched protein found to be reduced in PD441 was VWA3B. VWA3B is expressed in respiratory epithelium and thought to function in DNA transcription and repair. It was predicted to be a ciliary protein in the CiliaCarta (Van Dam et al., 2019) and CilDB databases (Arnaiz et al., 2009). VWAB3 contains the same transcription factor binding site as DNAH9, DNAI1, and CCDC151 (Patir et al., 2020). In the STRING interaction network, VWA3B appeared to interact with several HCs: DNAH9 and DNAH10 (Figure 3c). Lastly, DTHD1 is a protein containing the death domain and is likely involved in inflammatory signaling and programmed cell death (Hyun et al., 2007). Like KIAA1430 and VWA3B, DTHD1 was reduced in the DNAH5-defective samples. DTHD1 has been found to interact with CCDC65 (Figure 3c), a component of the N-DRC associated with PCD (Horani et al., 2013; Jreijiri et al., 2024).

### Study limitations and future directions

It is important to note that our results are limited by sample size, resolution, and sensitivity. Our proteomics results indicate the presence of proteins previously unknown as related to respiratory motile cilia, namely CFAP97, VWA3B, and DTHD1 in wild type samples and their absence in patient samples. However, challenges in genetic engineering and limited resolution in cryo-ET imaging mean that the exact location of these proteins within the axoneme are difficult to determine. Additionally, it is unclear whether these proteins are stable components of the axoneme or part of more transient processes, such as IFT. Further investigations are needed to determine if and how these proteins function within the cilium.

Limited other cases of MS use in PCD research are known. Previous research used MS to identify inner dynein heavy chain DNAH7 as a missing protein in PCD axonemes in a targeted way (Zhang et al., 2002). Additionally, researchers have used a MS-based approach to broadly characterize bronchoalveolar lavage fluid exosomes from PCD and cystic fibrosis patients (Rollet-Cohen et al., 2018). To our knowledge, our report constitutes the first comprehensive, untargeted study of protein changes in isolated PCD axonemes from ALI-cultured cells. Whether characterization of axonemal composition by MS will lead to a better understanding of the genotype-phenotype association in patients with PCD needs to be examined in larger patient cohorts.

### Conclusion

Overall, by comparing isolated cilia from normal and PCD cells using traditional diagnostic methods, MS, and cryo-ET, we highlighted the similarities and differences in motile ciliopathy phenotypes. Our findings contribute to the understanding of how distinctive mutations in ODA genes, specifically truncations, affect axonemal composition and structure. We present the first use of MS to probe global changes in axonemal protein composition in PCD samples as a result of specific PCD gene variants. The precise role that MS and high-resolution imaging can play in the analysis of such patient material will need to be investigated in larger patient studies.

## Acknowledgements

We thank ScopeM and CEMK for technical assistance with electron microscopy. This work was financially supported by PHRT(#2021-365) and SNF Bilateral grant (#IZLIZ3_200294) to T.I.

## Abbreviations

ALI: Air-liquid interface
CBF: Ciliary beating frequency
CBP: Ciliary beating pattern
Cryo-ET: Cryo-electron tomography
ERS: European Respiratory Society
HC: Heavy chain
HSVM: High speed video microscopy
IDA: Inner dynein arm
IF: Immunofluorescence
ODA: Outer dynein arm
MTD: Microtubule doublet
NEC: Nasal epithelial cell
PCD: Primary ciliary dyskinesia
LC-MS: Liquid chromatography coupled to mass spectrometry
RS: Radial spoke
RT: Room temperature
STA: Subtomogram averaging
TEM: Transmission electron microscopy

## Appendix

**Figure S1.**
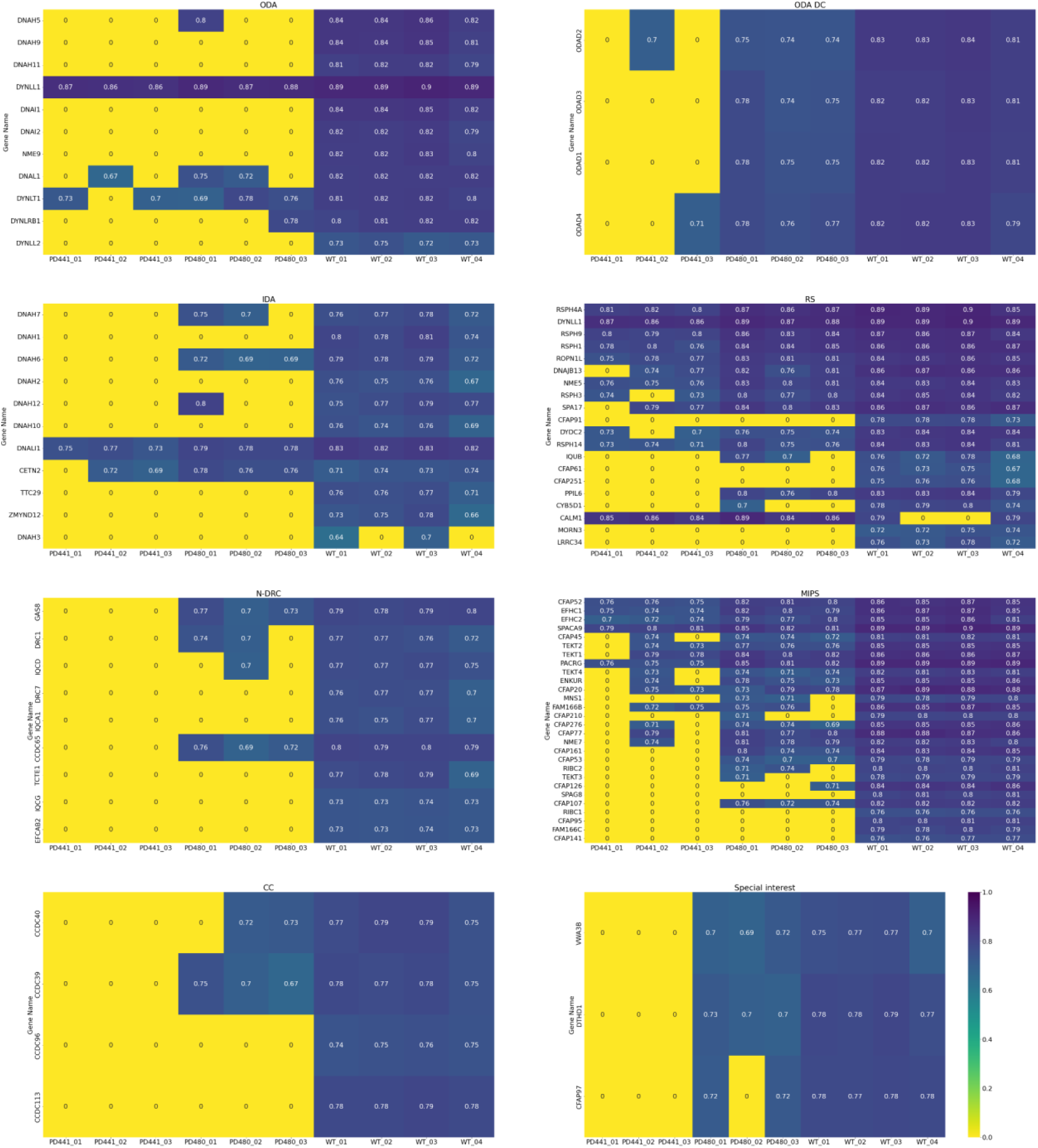
Heat maps summarizing the LC/MS results for PD441, PD480 and wild type samples.

**Figure S2.**
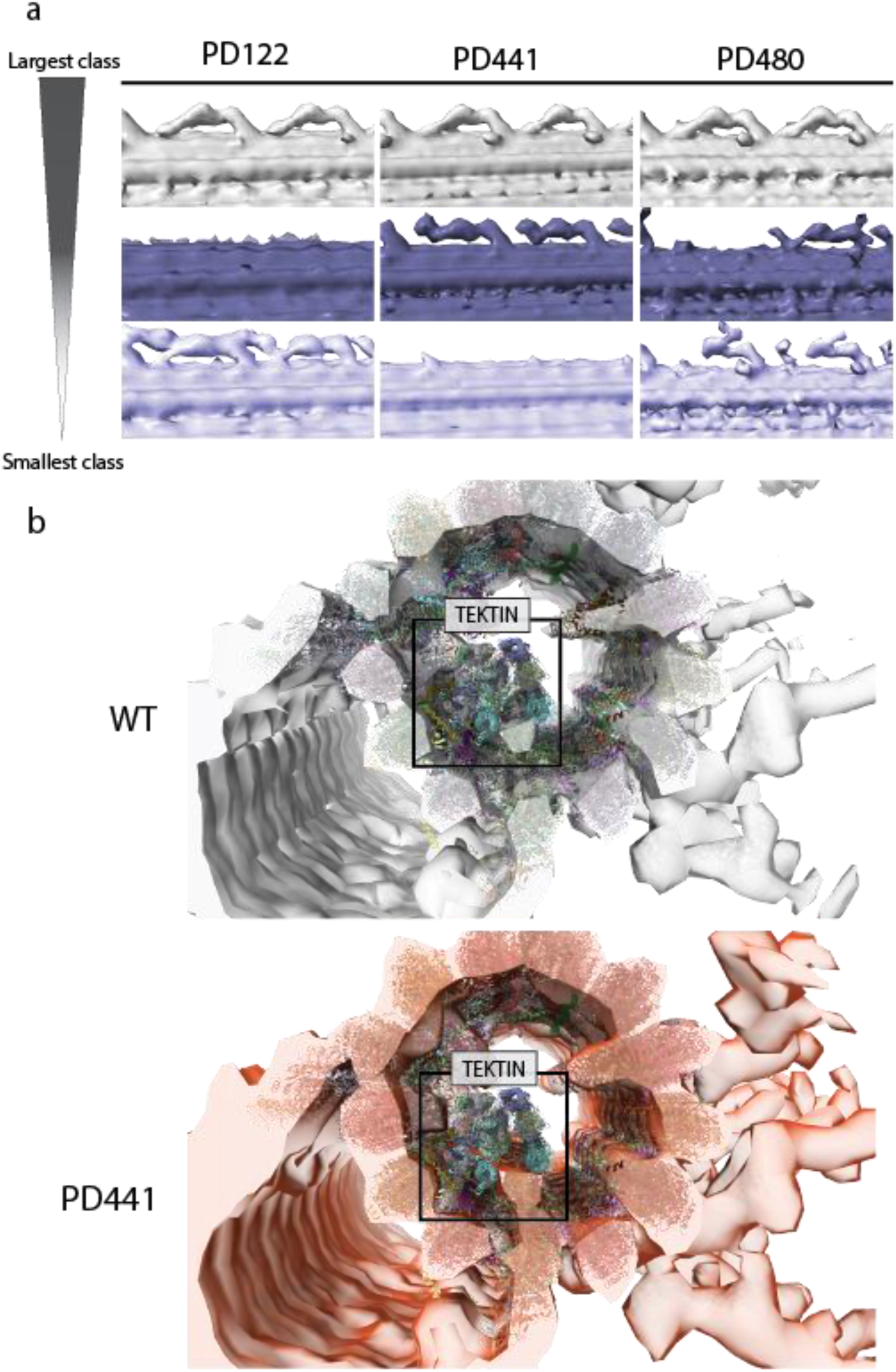
Structural analysis of PCD patient 24- and 96-nm periodic unit of the microtubule doublet (MTD) A. Classification of ODA particles for each patient. PD122 and PD441 show classes that have a complete loss of the docking complex, whereas all PD480 classes have B. Fitting of atomic model 7RRO [Gui et al 2021] into the 96-nm modular repeat tomographic maps of WT and PD441. Images generated in ChimeraX (Meng et al., 2023).

**Table S1.**
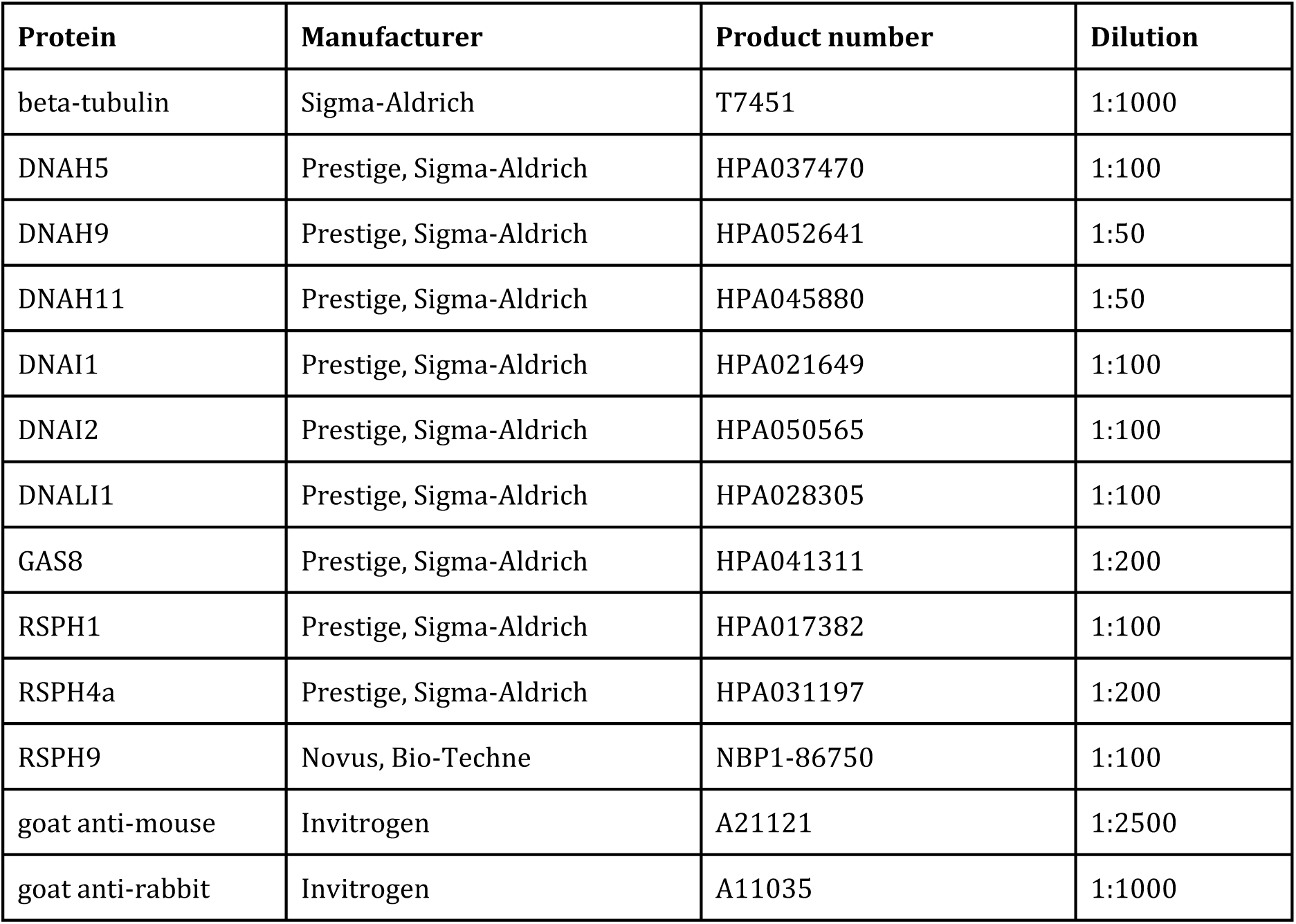
Antibodies for IF staining.

| Protein | Manufacturer | Product number | Dilution |
| --- | --- | --- | --- |
| beta-tubulin | Sigma-Aldrich | T7451 | 1:1000 |
| DNAH5 | Prestige, Sigma-Aldrich | HPA037470 | 1:100 |
| DNAH9 | Prestige, Sigma-Aldrich | HPA052641 | 1:50 |
| DNAH11 | Prestige, Sigma-Aldrich | HPA045880 | 1:50 |
| DNAI1 | Prestige, Sigma-Aldrich | HPA021649 | 1:100 |
| DNAI2 | Prestige, Sigma-Aldrich | HPA050565 | 1:100 |
| DNALI1 | Prestige, Sigma-Aldrich | HPA028305 | 1:100 |
| GAS8 | Prestige, Sigma-Aldrich | HPA041311 | 1:200 |
| RSPH1 | Prestige, Sigma-Aldrich | HPA017382 | 1:100 |
| RSPH4a | Prestige, Sigma-Aldrich | HPA031197 | 1:200 |
| RSPH9 | Novus, Bio-Techne | NBP1-86750 | 1:100 |
| goat anti-mouse | Invitrogen | A21121 | 1:2500 |
| goat anti-rabbit | Invitrogen | A11035 | 1:1000 |

**Table S2.**
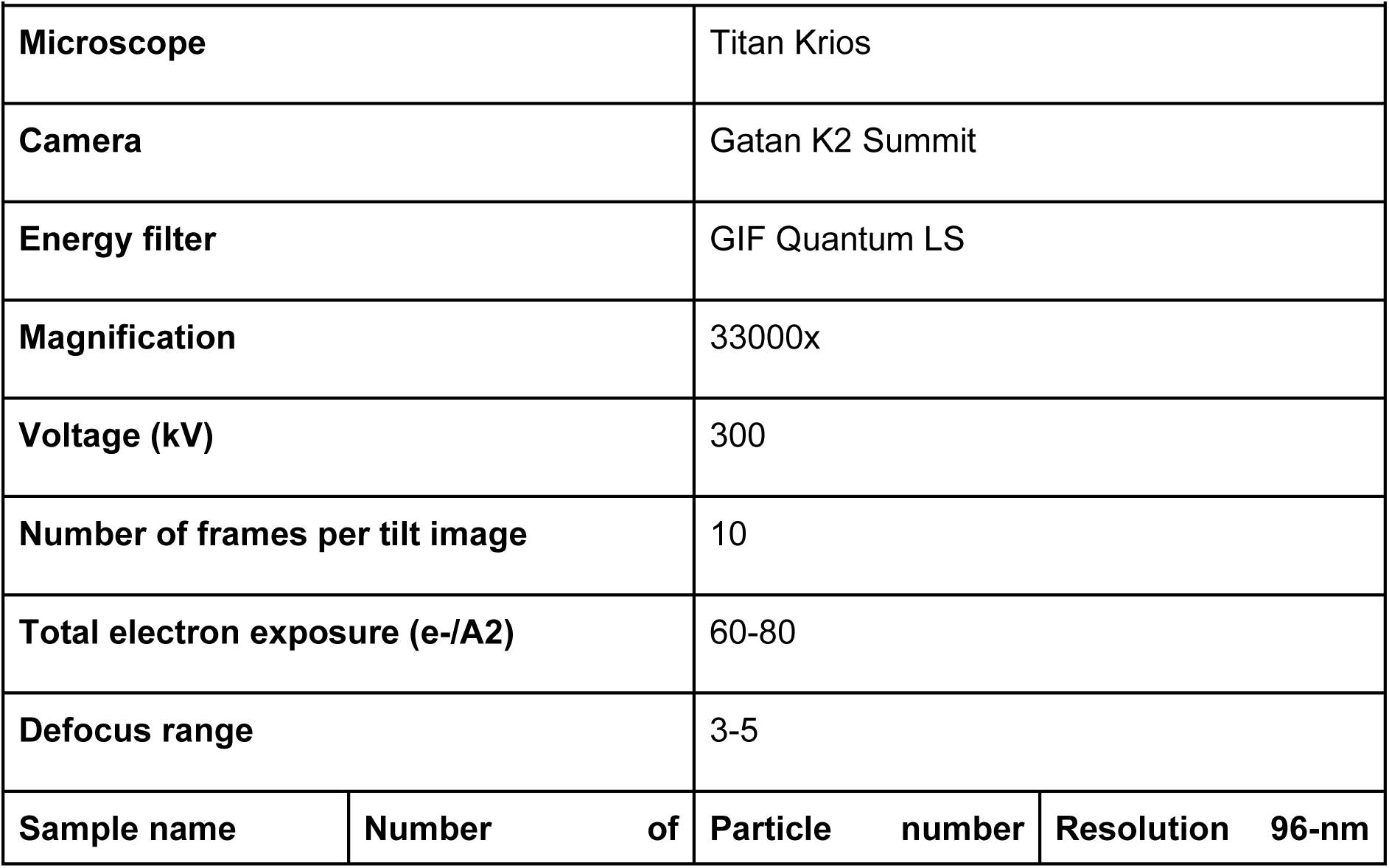
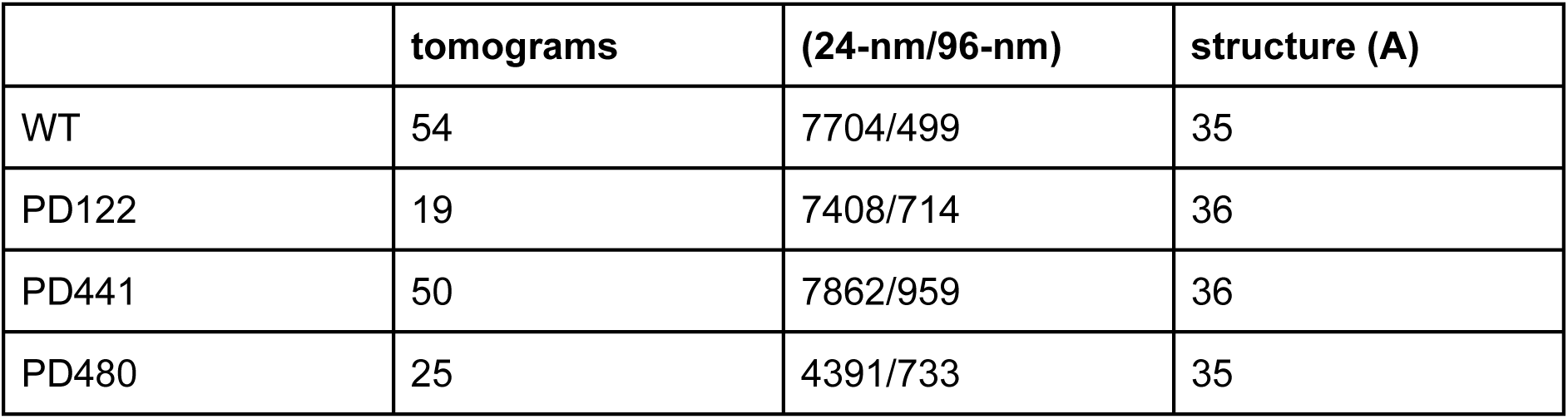
Tomography data analysis parameters for wild type and patient data.

**Table S3.**
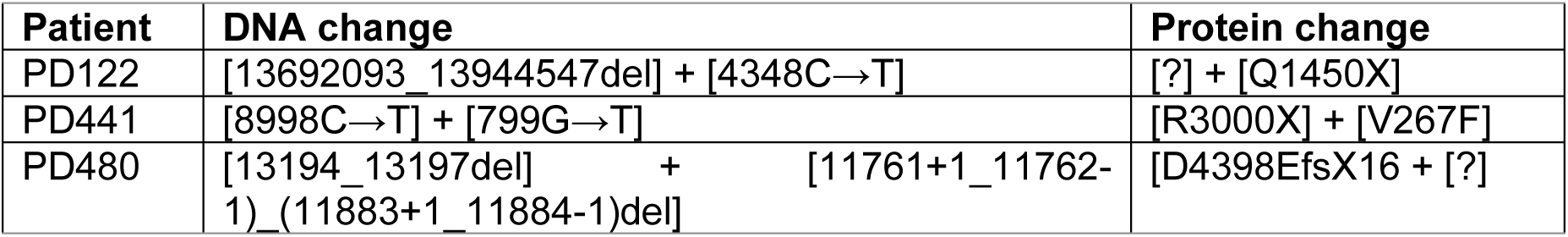
DNAH5 mutations of patient samples used in this study.

**Table S4.**
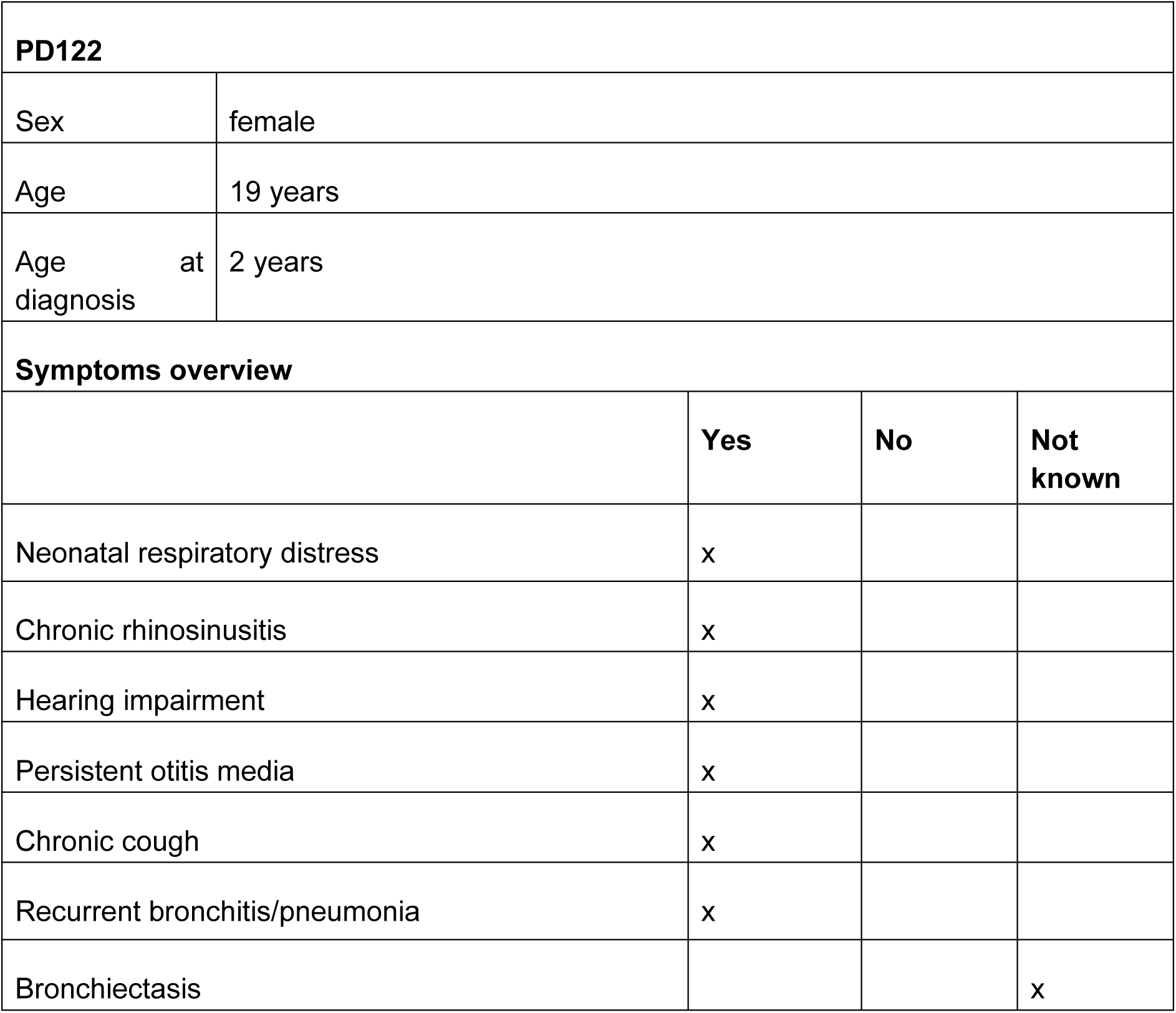
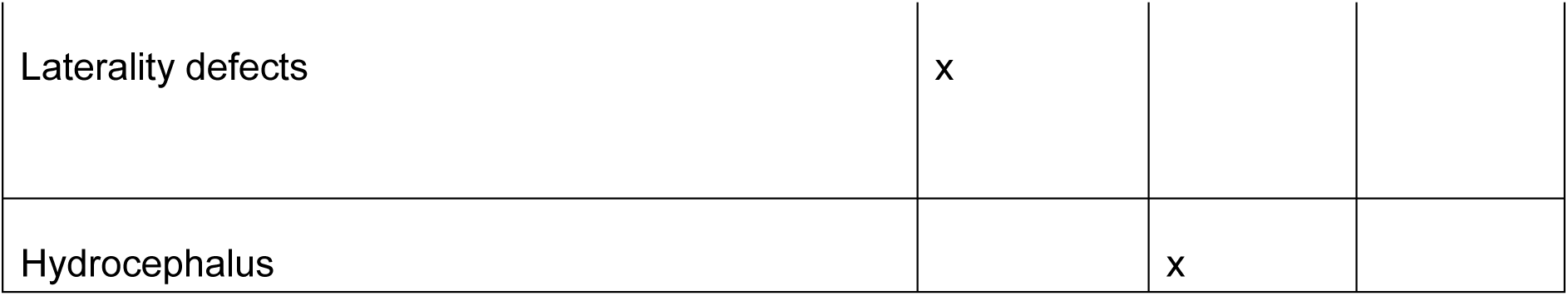
Clinical information concerning PD122.

|  |  |  |  |
| --- | --- | --- | --- |
| PD122 |  |  |  |
| Sex | female |  |  |
| Age | 19 years |  |  |
| Age at diagnosis | 2 years |  |  |
| Symptoms overview |  |  |  |
|  | Yes | No | Not known |
| Neonatal respiratory distress | x |  |  |
| Chronic rhinosinusitis | x |  |  |
| Hearing impairment | x |  |  |
| Persistent otitis media | x |  |  |
| Chronic cough | x |  |  |
| Recurrent bronchitis/pneumonia | x |  |  |
| Bronchiectasis |  |  | x |

|  |  |  |
| --- | --- | --- |
| Laterality defects | x |  |
| Hydrocephalus |  | x |

**Table S5.** Clinical information concerning PD441.

|  |  |  |  |
| --- | --- | --- | --- |
| PD441 |  |  |  |
| Sex | female |  |  |
| Age at brushing | 33 years |  |  |
| Age at diagnosis | 33 years |  |  |
| Symptoms overview |  |  |  |
|  | Yes | No | Not known |
| Neonatal respiratory distress | X |  |  |
| Chronic rhinosinusitis | X |  |  |
| Hearing impairment |  |  | x |
| Persistent otitis media |  | X |  |
| Chronic cough | X |  |  |
| Recurrent bronchitis/pneumonia | X |  |  |
| Bronchiectasis |  |  | x |
| Laterality defects |  | X |  |
| Hydrocephalus |  | x |  |

**Table S6.**
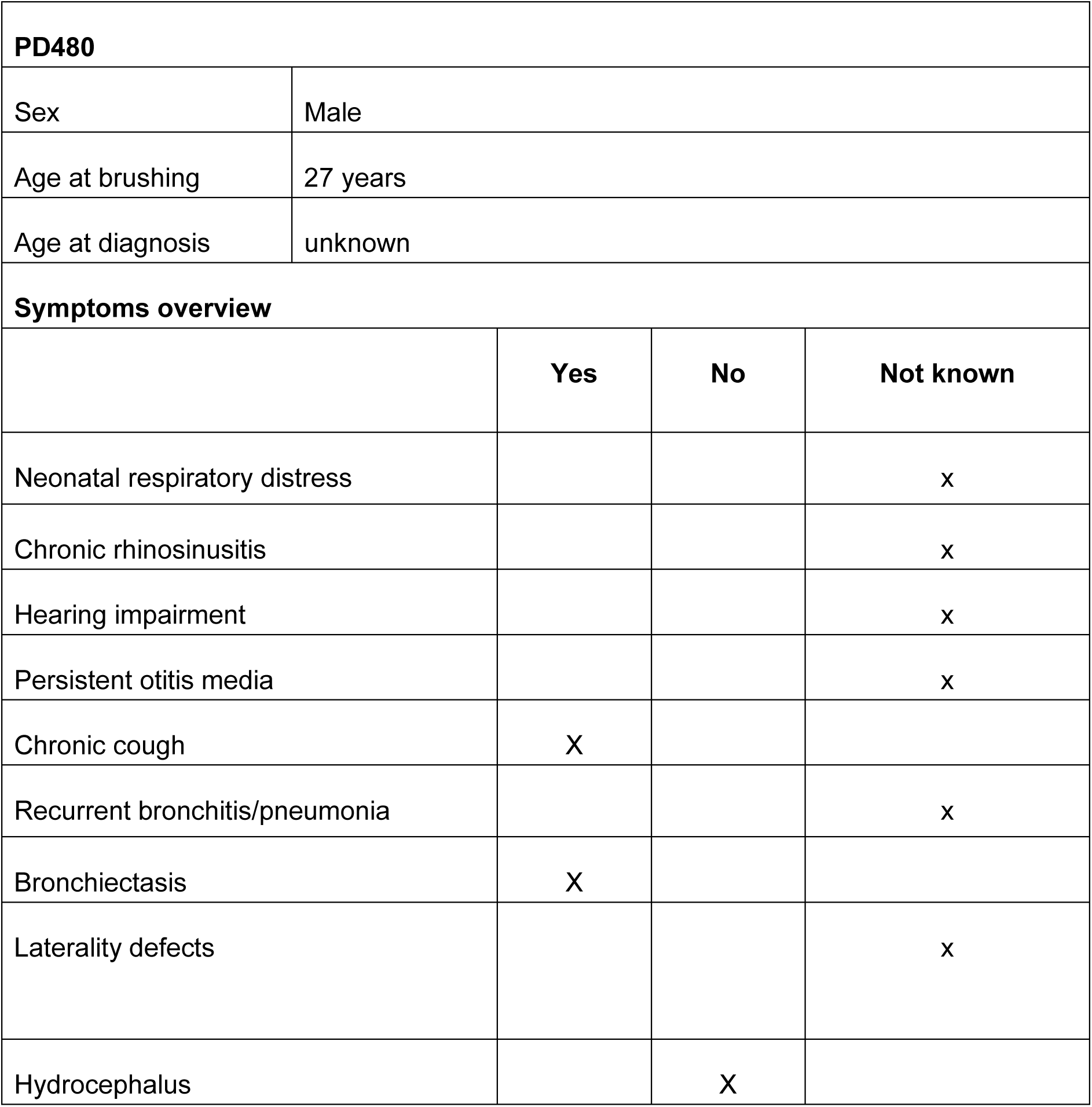
Clinical information concerning PD480.

